# Computer vision approach to characterize size and shape phenotypes of horticultural crops using high-throughput imagery

**DOI:** 10.1101/2020.07.24.199539

**Authors:** Samiul Haque, Edgar Lobaton, Natalie Nelson, G Craig Yencho, Kenneth V Pecota, Russell Mierop, Michael W Kudenov, Mike Boyette, Cranos M Williams

## Abstract

For many horticultural crops, variation in quality (e.g., shape and size) contribute significantly to the crop’s market value. Metrics characterizing less subjective harvest quantities (e.g., yield and total biomass) are routinely monitored. In contrast, metrics quantifying more subjective crop quality characteristics such as ideal size and shape remain difficult to characterize objectively at the production-scale due to the lack of modular technologies for high-throughput sensing and computation. Several horticultural crops are sent to packing facilities after having been harvested, where they are sorted into boxes and containers using high-throughput scanners. These scanners capture images of each fruit or vegetable being sorted and packed, but the images are typically used solely for sorting purposes and promptly discarded. With further analysis, these images could offer unparalleled insight on how crop quality metrics vary at the industrial production-scale and provide further insight into how these characteristics translate to overall market value. At present, methods for extracting and quantifying quality characteristics of crops using images generated by existing industrial infrastructure have not been developed. Furthermore, prior studies that investigated horticultural crop quality metrics, specifically of size and shape, used a limited number of samples, did not incorporate deformed or non-marketable samples, and did not use images captured from high-throughput systems. In this work, using sweetpotato (SP) as a use case, we introduce a computer vision algorithm for quantifying shape and size characteristics in a high-throughput manner. This approach generates 3D model of SPs from two 2D images captured by an industrial sorter 90 degrees apart and extracts 3D shape features in a few hundred milliseconds. We applied the 3D reconstruction and feature extraction method to thousands of image samples to demonstrate how variations in shape features across sweetptoato cultivars can be quantified. We created a sweetpotato shape dataset containing sweetpotato images, extracted shape features, and qualitative shape types (U.S. No. 1 or Cull). We used this dataset to develop a neural network-based shape classifier that was able to predict Cull vs. U.S. No. 1 sweetpotato with 84.59% accuracy. In addition, using univariate Chi-squared tests and random forest, we identified the most important features for determining qualitative shape (U.S. No. 1 or Cull) of the sweetpotatoes. Our study serves as the first step towards enabling big data analytics for sweetpotato agriculture. The methodological framework is readily transferable to other horticultural crops, particularly those that are sorted using commercial imaging equipment.

## Introduction

The market value of a horticultural crop can be heavily dependent on its quality, particularly on physical characteristics such as size and shape. Consumers prefer produce that have specific shape properties [1–3], which are often referred in the literature as”Ideal,”“Grade A”, or defined by United State Department of Agriculture as “U.S. No. 1” [2, 4–6]. Despite having the same nutritional value as the ideally shaped produce, deformed or “Cull” products are often rejected by consumers. As a result, deformed crops can be a source of food waste [2, 7, 8] and significant financial loss to growers. This loss can be severe for crops with high shape variability (e.g., sweetpotatoes and bell peppers). With recent advancements in optical sorting technologies in the vegetable and fruit packaging industry and advancements in big data analytics, the quantification of shape and size characteristics at production scale could enable the identification of factors (i.e., environmental factors, genotype, and cultural practices) that contribute to shape deformation in horticultural crops. Through improved understanding of the underlying drivers of crop shape, growers could revise their cultural practices to promote crop consistency, leading to increased grower profits and reduced food waste. A major obstacle, however, to implementing big data analytics in support of crop quality assessment is the absence of efficient, high-throughput methods to quantify 3D features associated with crop shape. Many horticultural crops are regularly analyzed at packing facilities using high-throughput imaging equipment, but images captured at these facilities are exclusively used to sort fruits and vegetables into shipping boxes and containers, and the images are not stored or used for further downstream analyses. To date, no methodological framework exist for analyzing size and shape quality metrics from images collected from commercial sorting systems, leaving the images largely unused. Yet, with the proper technology, these images could be further scrutinized to log the size and shape characteristics of harvested crops at large production-scales. Though automated morphological feature extraction approaches have been proposed for several fruits and vegetables [1, 4, 9–17], these methods are neither transferable to industrial sorting facilities nor capable of generating large datasets due to their low throughput. Previously published methods have focused mostly on 2D morphological features (i.e., height, width, and aspect ratio) and are unsuitable for quantifying produce with highly irregular shapes (e.g., sweetpotatoes, bell peppers, cucumbers, and carrots). In addition, previous studies did not incorporate existing industrial imaging infrastructure, but instead designed or used independent systems for image acquisition, making the methods unsuitable to couple with existing industrial machinery [2, 4, 17, 18].

In this paper, we introduce a novel computer vision approach to extract 3D shape features from crop images and classify individual fruits and vegetables into grade classes. We use sweetpotato (SP), a highly variable and irregular crop, as a representative use case. We used digital images obtained from a commercially available sorter (capable of capturing 5 sweetpotato images per second per lane) installed at the Sweetpotato Breeding Program at the Horticulture Crop Research Station (HCRS) in Clinton, NC, to reconstruct three-dimensional models of sweetpotatoes. We calculated shape features from the 3D model that could not be extracted directly from 2D images (i.e., curvature, radii of cross-sections, and tail length). We applied the 3D reconstruction and feature extraction method to 12,579 image samples collected from a sweetpotato yield trial to demonstrate how variations in shape features across sweetptoato cultivars can be quantified. We created a sweetpotato shape dataset containing sweetpotato images, extracted shape features, and labeled qualitative shape types (U.S. No. 1 or Cull) for 1,332 of the 12,579 sweetpotatoes. We used this dataset to identify a machine learning architecture to best classify shape type. We found that a neural network classifier performed best, predicting Cull vs. U.S. No. 1 sweetpotato with 84.59% accuracy. In addition, using univariate Chi-squared tests and random forest, we identified curvature, length-width ratio, cross-sectional roundness, and cross-sectional diameters to be the most important features for determining qualitative shape (U.S. No. 1 or Cull) of sweetpotato.

The 3D reconstruction and feature extraction method allows us to capture the variation in shape features extracted from thousands of sweetpotatoes, paving a way to apply big data analytics to understand sweetpotato shape variation. Our method makes use of currently discarded commercial imagery and provides data that could enable downstream analytics for quantifying and understanding shape variation across cultivars, and identifying the factors responsible for these variations. Thus, in addition to supporting research on industrial agricultural production dynamics, our method has the potential to support plant breeding programs by objectively providing phenotypic metrics beyond yield that can be incorporated into breeding and selection processes for the development of high-value cultivars. In addition, we demonstrate that the extracted features can be used to train and test automated machine learning models for classifying individual fruits and vegetables by grade. Automatic shape classification has two benefits. First, it enables researchers to understand what percentage of a particular cultivar is marketable (qualitatively good). Second, in the context of sweetpotato specifically, existing industrial sorters do not effectively capture sweetpotato shape features and fail to accurately sort SPs based on shape in an automated way. Due to the ability to calculate 3D features in milliseconds, our method can be incorporated into existing industrial sorters to improve their performance. Industrial deployment of this method will help packers improve accuracy and efficiency of the existing grading process (by reducing manual labor), and will also create novel datasets that can be used to analyze industrial-scale trends in crop quality.

## Materials and Methods

We developed a computer vision algorithm for creating 3D sweetpotato models from images captured by the Exeter Accuvision Sorter (Exeter Engineering, Exeter, CA). We extracted *thirteen* 3D shape features from the 3D SP model. We performed validation experiments to evaluate the accuracy of our 3D modeling and feature extraction method. We applied our feature extraction method to extract shape features from 12,579 SP images and quantified the distributions of these features across different cultivars. In addition, using a labeled dataset of 1,323 SPs and we trained and validated machine learning classifiers for identifying U.S. No. 1 vs. Cull SPs. Using Chi-squared test and random forest analysis we identified the influential features that determined SP shape class. Finally, by evaluating multiple performance metrics, we selected the champion classification model for SP shape type prediction. Fig 1 represents an overview of the methodology.

**Fig 1.**
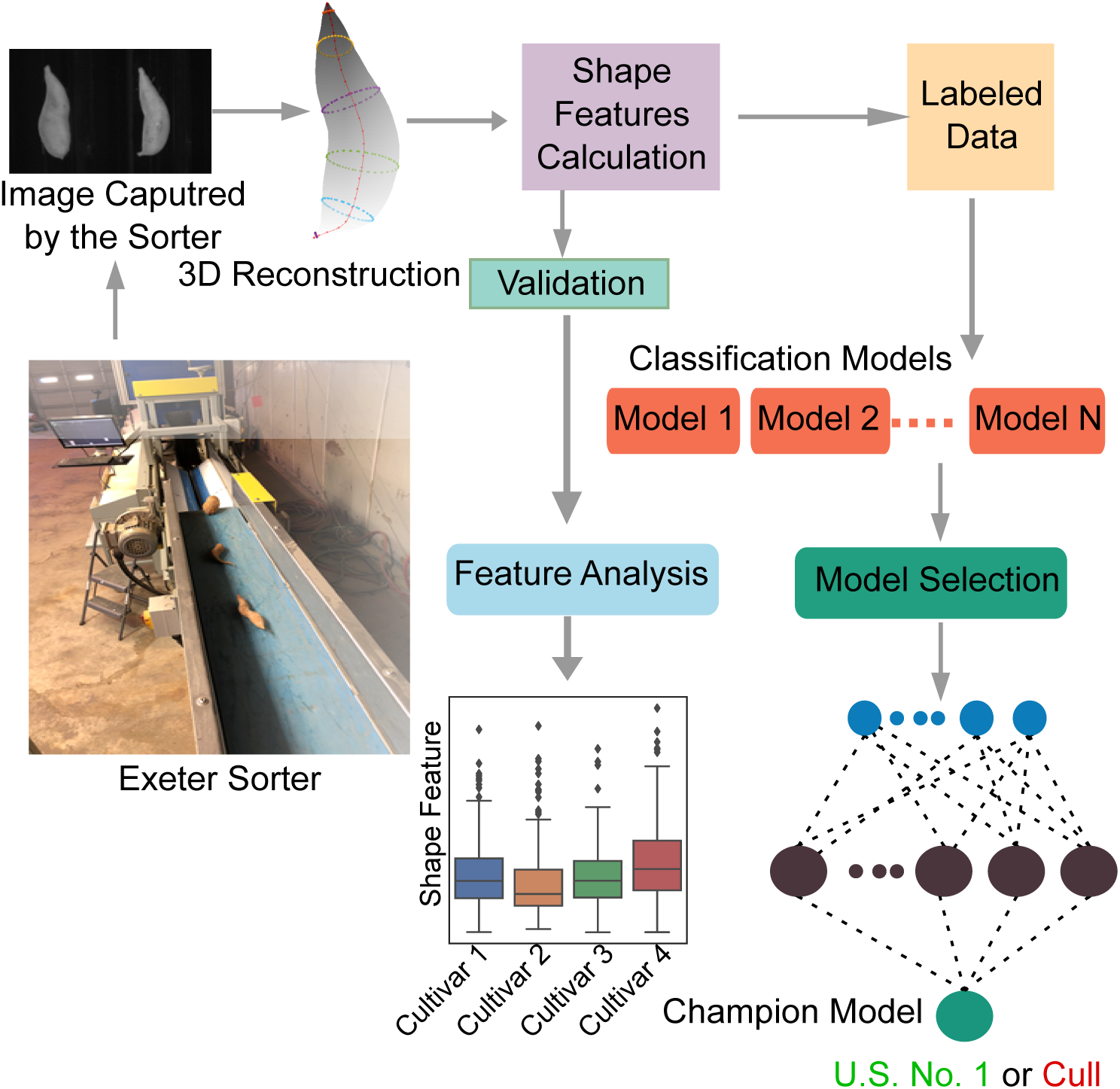
SP shape feature quantification and shape classifications. SP images were captured using a single lane sorter developed by Exeter Engineering, installed at the Horticulture Crop Research Station (HCRS) in Clinton, NC.

### Industrial Packing and Imaging of Sweetpotatoes

We obtained 12,579 sweetpotato images captured by an Exeter Accuvision Sorter installed at the North Carolina Department of Agriculture and Consumer Services (NCDA&CS) Horticultural Crop Research Station in Clinton, NC. The Exeter Accuvision Sorter can scan tens of thousands of SPs per hour and captures images of all SPs processed through it (Fig 1). This equipment is currently used by many packers for sorting different fruits and vegetables (including sweetpotatoes). We used a sorter with a single lane, whereas industrial packers use the same sorter with multiple lanes. The Exeter sorter captures Near Infrared (NIR) and Color (RGB) images of sweetpotatoes. Both images contain SP views from two separate angles that are 90*°* apart from each other. We used the NIR images for image processing (Fig 2A) and 3D reconstruction as these images are less noisy than the RGB images.

**Fig 2.**
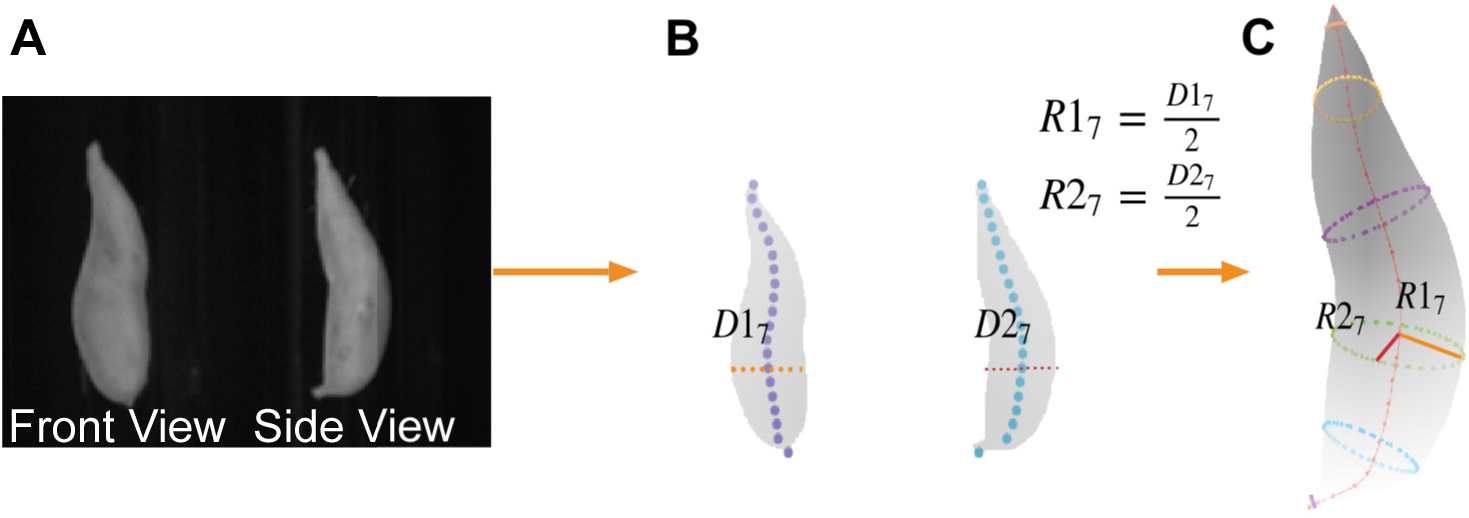
Sweetpotato 3D reconstruction. A) NIR image from the Exeter Accuvision Sorter, B) Segmented sweetpotato from the NIR images, central axes is obtained for each view, diameters are obtained across the axial length, C) Ellipsoidal reconstruction using the radii obtained from segmented sweetpotato image.

### 3D Reconstruction

We segmented sweetpotatoes from the NIR images (Fig 2A) using intensity-based thresholding. The segmentation provided sweetpotato shape outlines viewed from two different angles normal (i.e., 90*°* apart) to each other (front view and side view). We aligned and rotated the segmented sweetpotato images and calculated centroid axes for each segmentation mask. We selected *N* equidistant points across the axes and obtained sweetpotato radii at these points for both views, giving us *N* pairs of radii. Next, in a new 3D coordinate system, we constructed *N* ellipses on the *XY* plane along the Z-axis using the radii pairs. These ellipses were interpolated across the Z-axis to obtain reconstructed 3D SP shape. We implemented the 3D reconstruction methods and all image processing tasks in MATLAB R2020a (MathWorks, Natick, MA). We set the number of equidistant points *N* to 20. The number of points can be changed if necessary (based on the size of fruit or vegetable). However, increasing *N* will increase the number of computations needed for the 3D reconstruction.

### Camera Scale Factor Calculation

We designed an experiment to calculate the camera scale factor and assign specific units to the 3D model. We used a model sweetpotato with known height and width and scanned it using the Exeter scanner to obtain the model sweetpotato’s NIR images. We scanned the model SP nine times at different orientations and estimated the 3D shape for all the scans. Next, we used the known measurements to calculate the camera calibration factor for each scan. The average calibration factor obtained from these nine scans was used to calculate measurements for all other scans. Details of how the scale factor was calculated are provided in S1 Text.

### Shape Features

We used the reconstructed 3D model to calculate 13 SP shape features (Fig 5). Among these features, two (cross-section diameter, cross-section roundness) were calculated across 31 cross-sections of sweetpotato, giving us 73 shape variables in total. Cross-sections were normal to the curved SP axes. We used 31 cross-sections to ensure we had enough cross-sectional information for all sweetpotatoes in our dataset (adjacent cross-section centroids were 0.47 inches apart for the longest SP). The number of cross-sections is an arbitrary parameter and can be changed as needed. Primary shape features include curved length, straight length, maximum diameter/width, and diameter across cross-sections of the SP. In addition, we calculated several secondary shape features using these primary features. We calculated tail length by incorporating cross-sections from the edges of SP that have a diameter less than or equal to 1.5 inches. Curvature was calculated by taking the ratio of curved length to straight length. We calculated the length to width ratio using straight length and maximum diameter. We also calculated the ratio between tail length and body length. A complete list of extracted shape features is presented in Table 1. Details of feature calculation are available in S2 Text. We extracted shape features for all the available SP samples. We quantified the distributions of different shape features across SP cultivars.

**Table 1.** Shape Features. Calculated shape features are listed in the table. *indicates features calculated at every SP cross-section.

| Feature Name | Description |
| --- | --- |
| Axial / Curved Length ($L_C$) | Length across the central axis of SP |
| Straight length ($L_S$) | Tip to tip length of sweetpotato |
| Maximum diameter / width (W) | Estimated maximum diameter or width |
| Tail length ($L_{Tail}$) | Tail length estimated by calculating tip areas with a diameter less than 0.5 inch |
| Body length (without tail) $L_B$ | $L_C - L_{Tail}$ |
| Length to width ratio | $\frac{L_S}{W}$ |
| Curvature | Curvature is calculated as the ratio of curved length to straight length, with adjustment for tail length. $\frac{L_C - L_{Tail}}{L_S - L_{Tail}}$ |
| Diameters across cross-sections* | Diameter / Width of each cross-section |
| Tail to axial length ratio | $\frac{L_{Tail}}{L_C}$ |
| Tail to body length ratio | $\frac{L_{Tail}}{L_B}$ |
| Volume (V) | Volume estimated from $L_c$ and cross-sectional areas |
| Cross-section roundness* | Standard deviation of distances from cross-section center to perimeter normalized by diameter of cross-section. |
| Average Cross-section roundness | Mean of the standard deviations of radii across cross-sections. |

### SP Weight Estimation

Our approach can also be used to estimate individual SP weight from SP density and estimated volume. We estimated the volume of an individual SP using features extracted from the 3D model (S2 Text). To estimate SP density, we measured weights of 19 randomly sampled sweetpotatoes of varying shape and size. Then we scanned each sweetpotato multiple times (16 SPs were scanned 4 times and 3 were scanned 5 times giving us 79 images) using the Exeter sorter. We estimated the density for each scan by using the known weight to obtain the mass and dividing the mass by the estimated volume 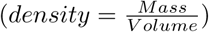. We used the average density value to estimate weights for individual sweetpotatoes.

### Shape Classification

We used a machine learning-based shape classifier to asses the extent to which our extracted features could be used to discriminate between marketable and unmarketable SPs. To train a machine learning-based shape classifier that takes extracted shape features as input and generates predicted shape label as output, we created a database of 1,332 labeled or classified SP images scanned using the Exeter Accuvision Sorter. Each image was labeled by a domain expert (researcher from the NC State University sweetpotato breeding program) as either U.S. No. 1 (a sweetpotato that will have high market value and meets the U.S. No. 1 standard established by USDA Agricultural Marketing Service [5]) or Cull (a sweetpotato that will potentially have a lower market value or will be discarded during harvesting/sorting) based on its visual properties. In addition, we categorized the Cull sweetpotatoes into four qualitative shape classes: Tailed, Tapered, Curved, and Other (Fig 3). We then extracted shape features for all labeled images. We partitioned the labeled dataset into 80% training and 20% holdout (used for evaluating classification performance) set. This gave us 1066 (494 U.S. No. 1 SP and 572 Culls) training samples and 266 (123 U.S. No. 1 and 143 Cull) holdout samples. Using the extracted features and assigned labels from the training set, we trained binary classifiers (for classifying U.S. No. 1 and Cull SPs) models using SAS Viya V03.05 Model Studio (SAS Institute Inc., Cary, NC). Multiple machine learning models (decision tree [19], neural network [20], random forest [19], logistic regression [21, 22], Bayesian network [22] and gradient boosting [19]) were trained and tuned (hyperparameter selection) using a 5 fold cross-validation of the training data set (70% training and 30% validation in each fold). We selected the champion model by comparing different performance metrics (Accuracy, F1 Score [23], and Area Under Receiver Operator Characteristics [AUROC] curve [24]) of all the trained classification models.

**Fig 3.**
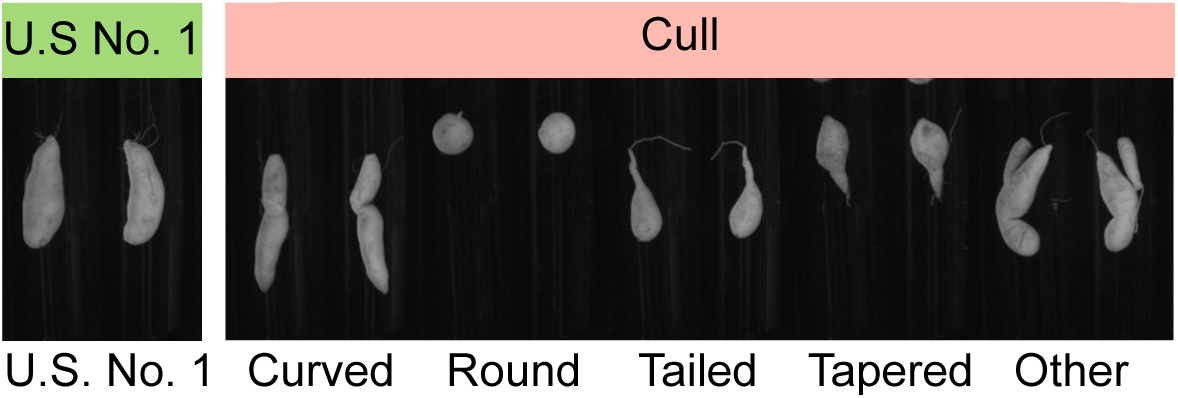
Sweetpotatoes with different shape types. Images captured using Exeter Accuvision Sorter.

### Variable Importance Analysis

To understand which shape features played influential roles in determining shape label (U.S. No. 1 or Cull), we conducted a variable importance analysis. We used the Chi-squared test [25–27] (using MATLAB’s *fscchi2* function) to examine the dependency between shape class and each shape feature. The random forest classifier [19] also produced a ranking of important variables based on the change of the residual sum of squares [28]. Variable rankings from these two methods provided insight into important features for shape class determination.

## Results

### 3D Reconstruction of Sweetpotato

Using our 3D reconstruction approach, we generated 3D models for all 12,579 imaged sweetpotatoes. Fig 4 shows reconstructed models for SPs of various shape types. The MATLAB implementation of our approach produced reconstructed 3D model of a SP within a few hundred milliseconds (on an Intel Core i7 processor with 16 GB Memory). In its current implementation, this algorithm can be used to calculate SP shape features at production-scale with very little delay. The speed of the method can be further improved by utilizing parallel processing of multiple images. Thus, this method can potentially be used in a high-throughput industrial sorter to capture SP features at the time of sorting.

**Fig 4.**
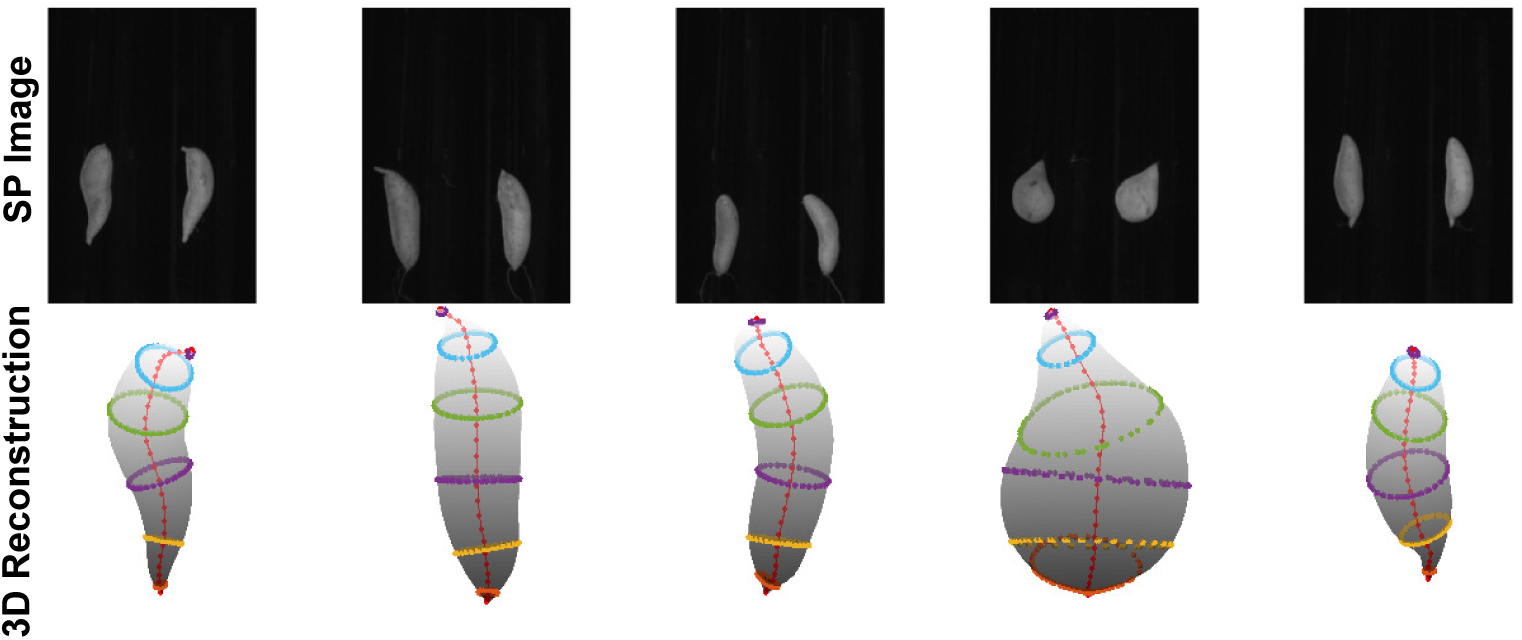
Sweetpotato 3D shapes. Examples of 3D reconstructed models of sweetpotato using images from Exeter Accuvision Sorter.

### Validation of Extracted Features

We validated the accuracy of extracted shape features (Fig 5) by measuring the length and maximum diameter (width) of randomly sampled SPs using slide caliper and measuring scale. We scanned these SPs using the Exeter sorter and estimated the same features using the 3D reconstructed model. Fig 6 shows that laboratory measurement and estimated measures are highly correlated (*R*^2^ = 0.958 and *R*^2^ = 0.923 for estimated straight length and maximum diameter, respectively). These results are strong indicators of the accuracy of the extracted features.

**Fig 5.**
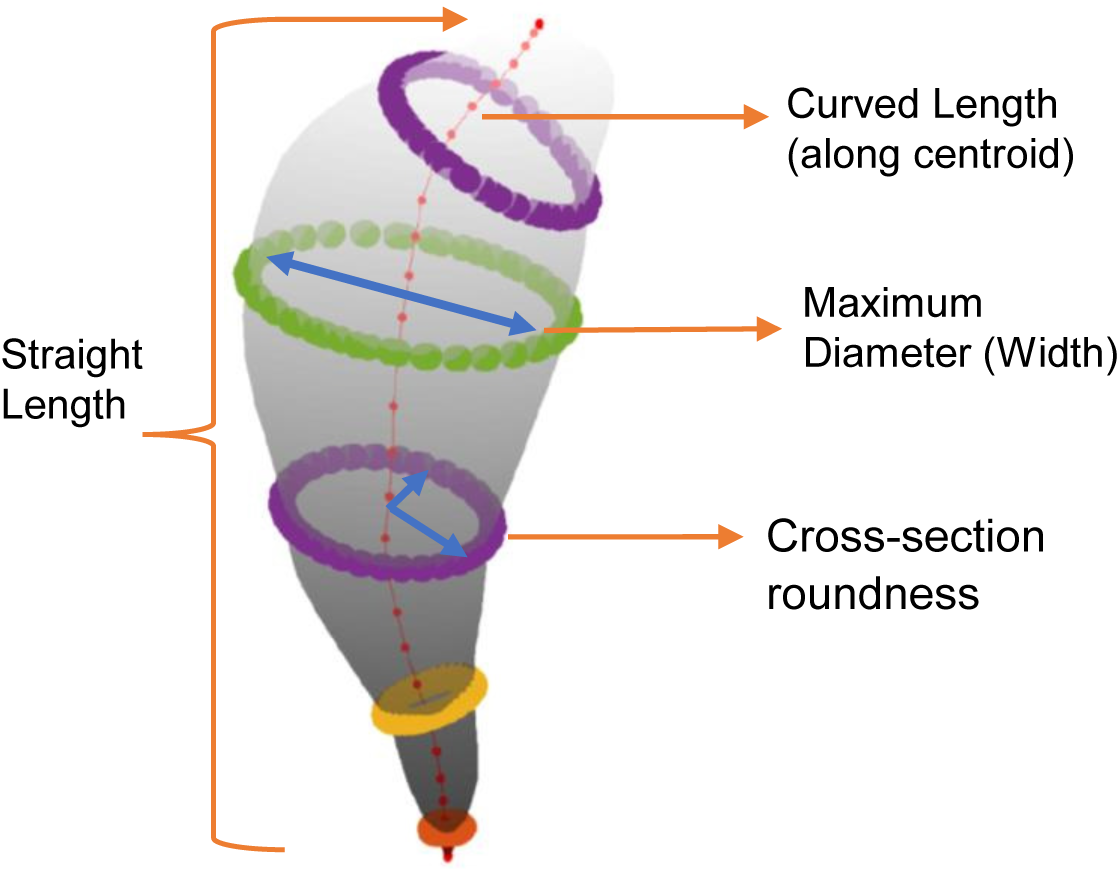
Sweetpotato Shape features. Extraction of different shape features from the reconstructed 3D model.

**Fig 6.**
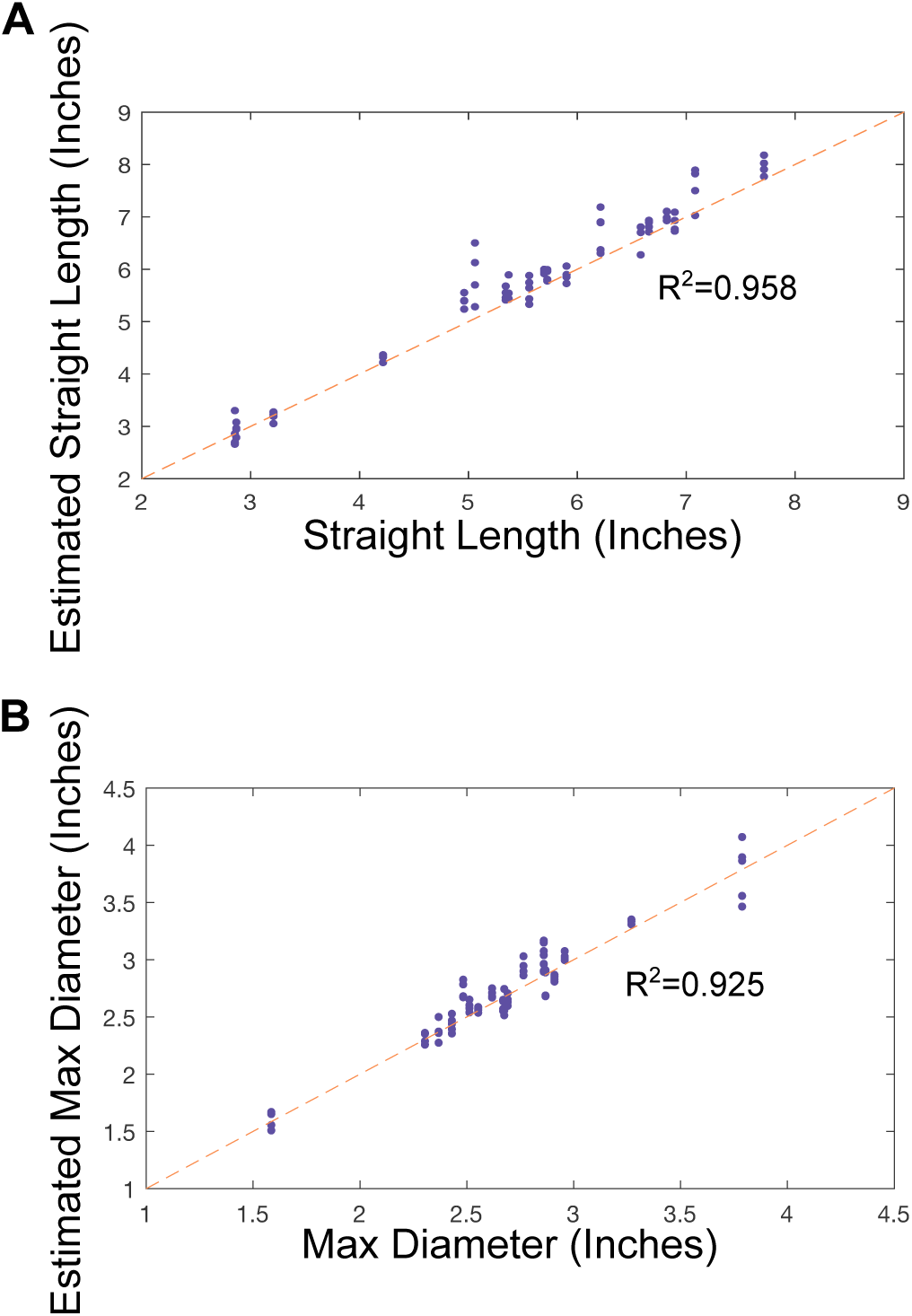
Scatter plots showing comparison between experimental measures vs estimated shape features in randomly sampled sweetpotatoes (n=79) (A) Estimated vs measured straight length (*R*^2^ = 0.958), (B) Estimated vs maximum diameter (*R*^2^ = 0.925).

### Application of Feature Extraction Algorithm

#### Shape Features Across Cultivars

The feature extraction algorithm enabled us to visualize the distribution of shape features across different cultivars. Fig 7 shows distributions of SP shape features in a subset of the data (1,943 sweetpotatoes, restricted to one field and four cultivars). For SP grown on that field, the Covington cultivar had the highest median width of 2.54 inches, while the Bellevue had the lowest median width of 1.95 inches. The distribution of curvature across all cultivars grown on that field was approximately the same with the Baeuregard cultivar having the smallest interquartile range.

**Fig 7.**
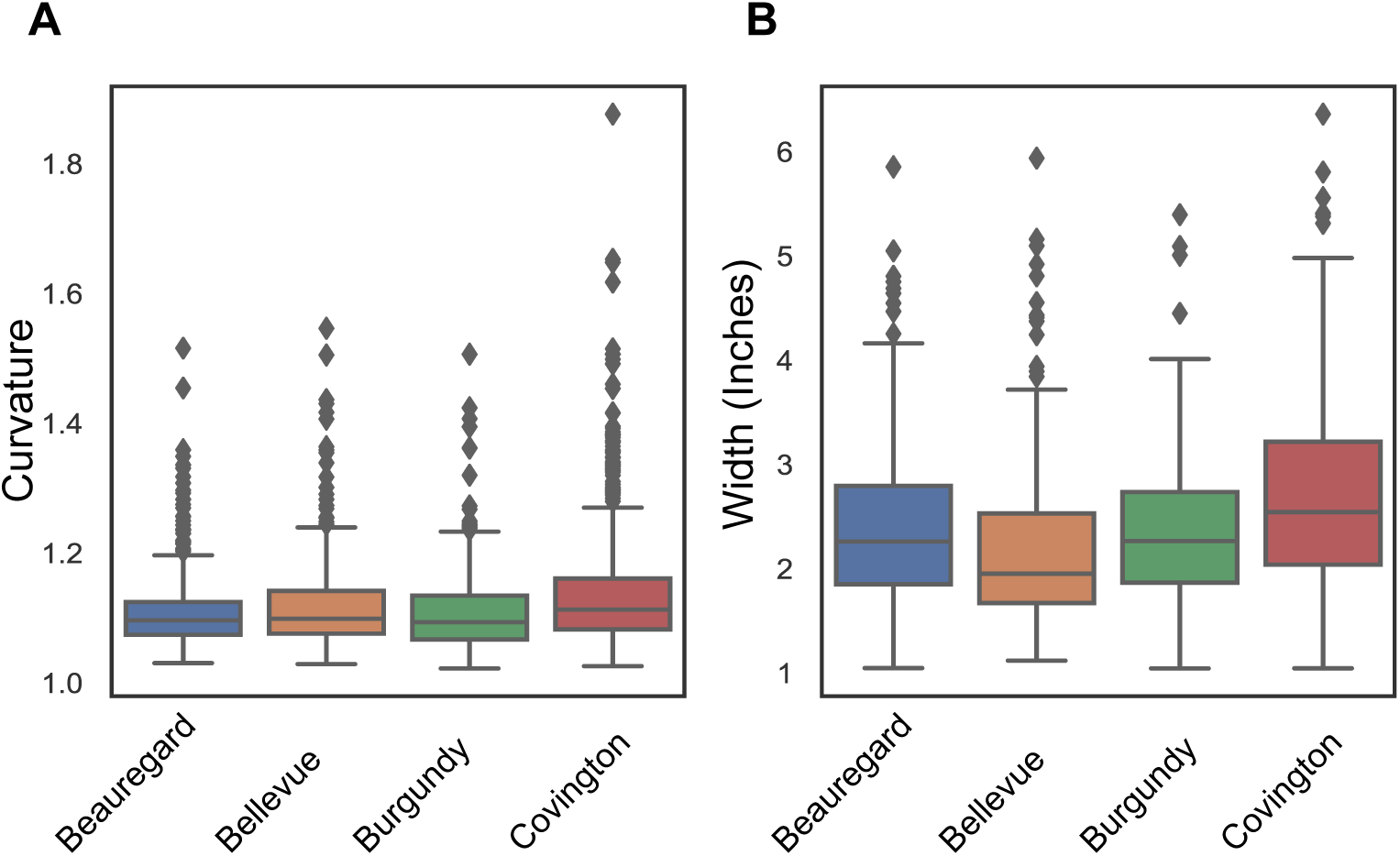
Box plots showing variation in shape features of sweetpotatoes sampled from a field trial in Clinton, NC. (A) Distribution of curvature across different cultivars, (B) Distribution of width across different cultivars.

#### SP Weight Estimation

We estimated the average density of 19 randomly sampled sweetpotatoes as 15.64 *grams/in*^3^ with a standard deviation of 1.27 *grams/in*^3^. Root mean squared error between the estimated weight and actual weight of the SPs was 27.4173 *grams*. We used the average density value to calculate the weights of 1,323 labeled SPs from their estimated volume. We want to point out that this is just an estimate of the density. Inaccuracies in this calculation could stem from unclean SPs that still contained soil on the surface and differences in density in bulk vs. tail parts of the SP. Fig 8 shows a plot of the variation in SP weight for SPs labeled as US No. 1’s vs SPs labels as Culls. The median estimated weight of the SPs labeled as U.S. No. 1 was 150.77 grams while the median estimated weight of the SPs labeled as Cull was 166.78 grams. Among the Cull SPs we found 6 SPs weighing above 1000 grams, 4 of these SPs belonged to the Other subclass, 1 belonged to the Round subclass, and 1 belonged to the Curved subclass.

**Fig 8.**
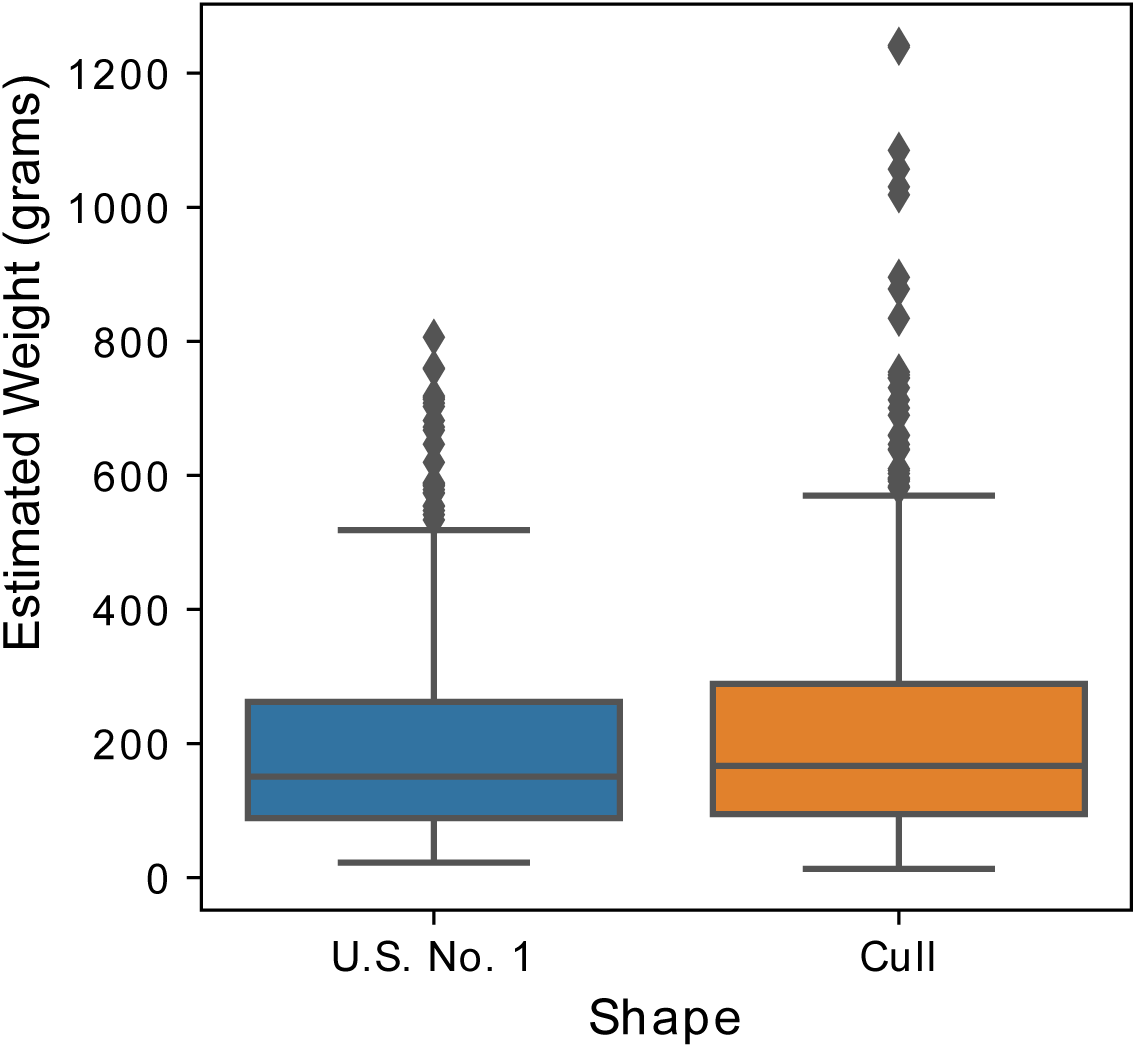
**Box plot showing variation in weights** among US. No.1 and Cull sweetpotatoes in our labeled dataset.

### Variable Importance

We analyzed relative importance of different shape features in determining U.S. No. 1 vs. Cull shape classes. The top 30 important features (for U.S. No. 1 vs Cull determination) identified by Chi-squared test [25–27] and random forest [28] are shown in Fig 9. We found that curvature and length-width ratio are the two most important features in determining shape labels and were identified by both methods. The rest of the common influential features are the roundness of cross-sections 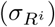 and diameters across cross-sections (*D*_*i*_). Random forest method also picked curved length, body length, and straight length as important factors for determining shape label.

**Fig 9.**
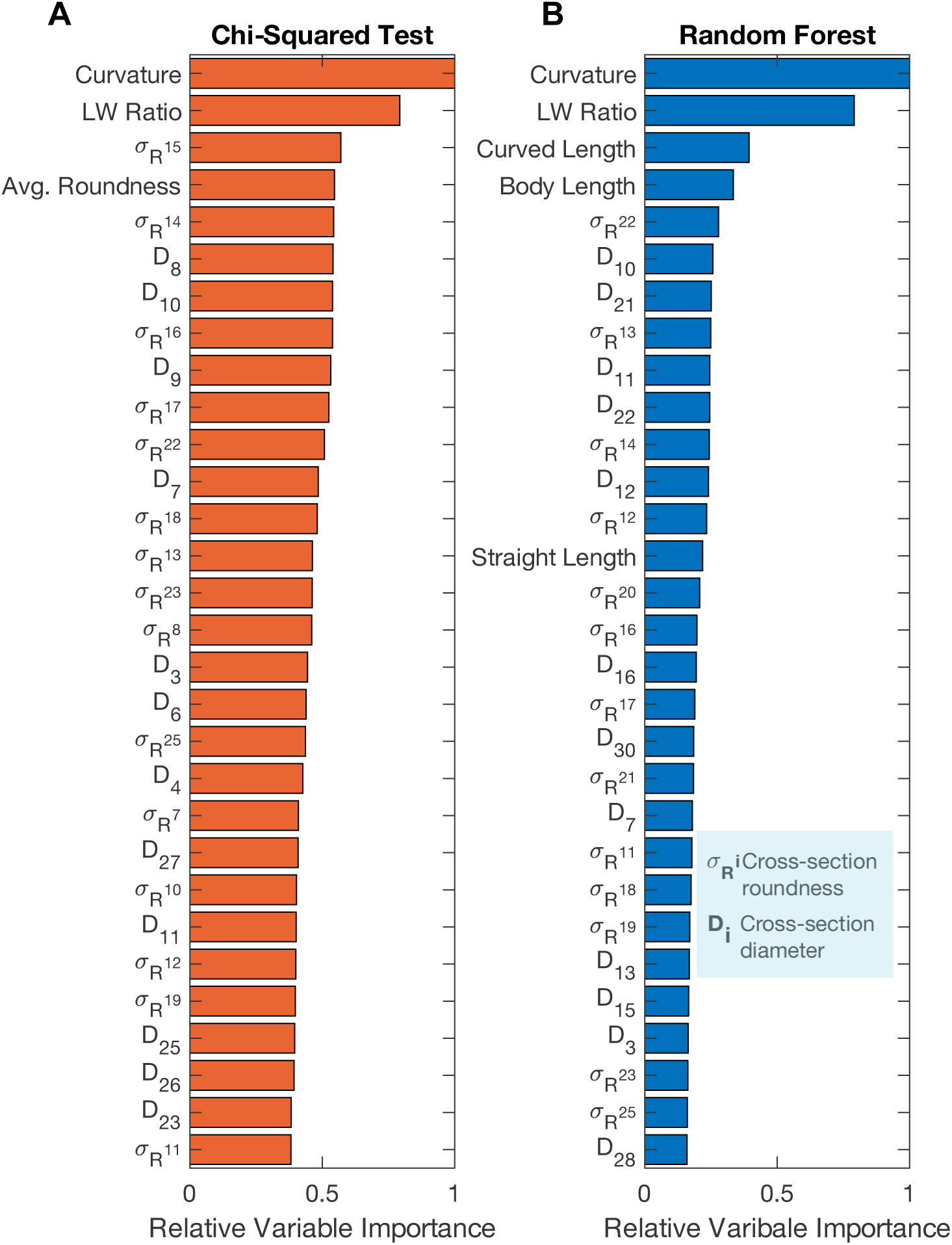
**Variable importance calculated using** (A) Chi-squared test, (B) Random forest. The X-axis represents relative variable importance scores. Curvature, length to width ratio (LW ratio), cross-section diameter (*D*_*i*_), and roundness 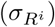 are identified as the most influencing features in determining shape labels by both approaches.

### Binary Shape Classification

Evaluation metrics for all competing classifier models are provided in Table 2. We obtained the optimum hyperparameters for the classifiers using the genetic search algorithm in SAS Viya (S4 Text). The neural network model yielded the highest accuracy and F1 Score [23] on the holdout set. The neural network had 1 hidden layer with 100 neurons with hyperbolic tangent activation function [20]. The confusion matrix [29] for the neural network model is given in Table 3. We obtained an accuracy 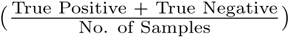 of 84.59%, F1 score [23] of 0.85, and AUROC [24] of 0.88 for the neural network model.

**Table 2.** Evaluation metrics (on the holdout data) for competing binary classification models. Neural network yielded highest accuracy and F1 score.

| Model | Accuracy | F1 Score | AUROC |
| --- | --- | --- | --- |
| Neural network | 84.59% | 0.848 | 0.88 |
| Gradient boosting | 83.08% | 0.830 | 0.857 |
| Logistic regression | 79.32% | 0.795 | 0.889 |
| Random forest | 78.20% | 0.783 | 0.827 |
| Decision tree | 75.56% | 0.754 | 0.772 |
| Bayesian network | 74.81% | 0.741 | 0.809 |

**Table 3.**
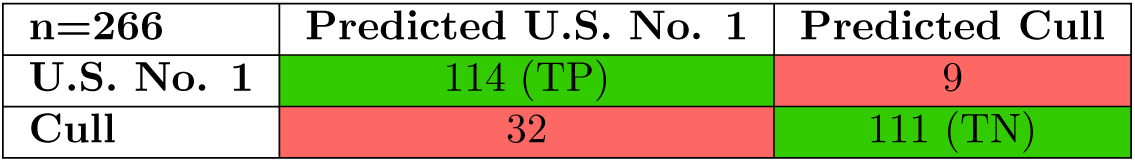
Confusion matrix for neural network classification, evaluated on holdout data. TP stands for True Positive and TN stands for True Negative. The overall accuracy on the holdout set is 84.59%

| n=266 | Predicted U.S. No. 1 | Predicted Cull |
| --- | --- | --- |
| U.S. No. 1 | 114 (TP) | 9 |
| Cull | 32 | 111 (TN) |

### Multi-class Classification

Our main goal was to identify Cull and U.S. No. 1 sweetpotatoes using the best performing classification model. However, we also wanted to assess the ability to detect subclasses of Cull sweetpotatoes using a machine learning model. We performed a model comparison analysis for the multi-class classification problem (S5 Table). The gradient boosting [19] model did better than the neural network [20] for multi-class classification. However, the overall accuracy of the gradient boosting multi-class classifier was only 65% on the holdout set. Gradient boosting model for multi-class classification achieved 91.87% sensitivity 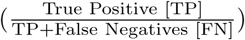 in predicting U.S. No. 1 sweetpotatoes and 80.76% sensitivity in predicting Round sweetpotatoes. However, the sensitivity for other class labels were dramatically lower (52.94% for Curved, 22% for Tailed, 21% for Tapered, and 0% for Other). The multi-class model predicted majority of the Tapered and Tailed sweetpotatoes as U.S. No. 1.

## Discussion

We developed a method that can accurately capture shape features (Fig 6) by reconstructing the 3D model of a horticultural crop from 2D images acquired by a high-throughput commercial sorter (Fig 5). To our knowledge, our method is the first to utilize existing industrial imaging equipment. It is also able to extract shape features significantly faster than previously reported shape extraction methods for different fruits and vegetables [2, 13, 16–18, 30, 31]. Most research investigating size and shape traits of horticultural crops used lab-scale imaging equipment that are slower and cannot be adapted to production scale environments [17, 32]. Further, we considered both ideal and deformed sweetpotatoes making our approach capable of quantifying loss due to shape deformation, while many prior studies excluded sub-standard produce [4, 18].

By using images acquired from already-used industrial equipment, the algorithm presented here can be readily implemented in production agriculture to gather and analyze large scale data. To deploy our method at a production facility, the requirements would be a desktop computer, MATLAB license, and an interface to image data. It is also possible to port this algorithm into an open-source language (e.g., Python, C++), eliminating the need for additional software licensing. A parallel programming implementation of the algorithm can be done with some modification of the existing code. This would allow the algorithm to process multiple images simultaneously and further increase the throughput of our approach. One major challenge in the industrial deployment of this method will be the management of vast amounts of shape data that will be produced from processing hundreds of thousands of SPs per hour. One way to mitigate this challenge is by uploading the data in a cloud server at regular intervals.

Our method paves the way for investigating underlying factors responsible for shape variations in different cultivars. As shown in Fig 7, we can quantify shape feature distributions across cultivars in large datasets, which can assist breeders in evaluating genotype *×* environmental interaction more effectively and can lead to the identification of potential new cultivars in less time. Our main goal was to demonstrate that we can quantify shape features across cultivars using the proposed computer vision approach. Our results for 1,943 SPs grown on one field (Fig 7) suggests that on average Covington SPs have higher curvature and width than those of the Beauregard, Bellevue, and Burgundy SP varieties. Expanding this approach to statistically assess the entire SP yield trial containing 12,579 sweetpotato samples from 14 different cultivars and grown in two different fields would require additional detailed analysis that incorporates the experimental designs of the SP yield trials.

Through variable importance analysis, we identified several key features that characterize SP shape by using the Chi-squared test [25–27], and random forest (Fig 9). Previous studies identified the LW ratio captured from 2D images as the standard feature for quantifying shape variations in agricultural produce [2, 33]. However, LW ratio alone is inadequate for capturing sweetpotato shape variation [18]. Our results show that curvature, cross-section roundness, and cross-sectional diameters are influential factors for determining shape class. Multiple previous studies reported volume as important shape features for horticultural produce [3, 18, 34, 35]. Interestingly, we did not find volume and tail length among the top 30 influencing variables. This result suggests that cross-sectional features (roundness and diameter) capture the information encoded in volume. We think that the ability to extract features from arbitrary cross-sections makes our method applicable to other crops with varying shapes and sizes. One of the most important traits of a sweetpotato is its weight. Fig 8 shows how our method can be utilized to obtain weight for each SP by shape class. It is essential to state that our density estimation was not accurate since we did not clean the SPs before scanning and weighing. In addition, we did not incorporate density variation among different cultivars. Thus, the actual weights of the SPs may be different but proportional to our estimates. With an accurate SP density measurement, our method can quantify weight distribution of SPs across shape class and cultivars, which will allow growers to better predict yields and pack-out from fields.

We used the extracted features to train a neural network classifier to classify SPs into Cull and U.S. No. 1 classes with reasonable accuracy (84.59% on holdout data) (Table 3). Among previous works, Okayama et al. achieved 95.7% accuracy in classifying bell peppers into Grade A and Grade B using a neural network classifier. However, their study used four side views and one top view (total five images) of a bell paper to extract 2D shape features from individual views, whereas our method uses just two side views of a SP. With two side views (90*°* apart) of a bell pepper their study achieved less than 60% classification accuracy with the 2D features (by applying statistical thresholds to the features), significantly lower than our results. We believe that, the capability of extracting 3D features allowed our method to perform better with just two views. We think that our method is deployable in industrial packing facilities for improved (i.e., more accurate and faster) automated sorting. This method can also be used in SP yield studies to quantify the amount of deformed SPs across cultivars and obtain a better estimate of post-harvest losses.

Though, the binary classifier performed well (Table 3), the accuracy of the multi-class classification was low (65% on holdout data). The multi-class classifier predicted the majority of Tapered and Tailed sweetpotatoes as U.S. No. 1 (Table 4). The model struggled to learn multiple cull shape labels with available data. Overall accuracy on the training set was only 78%. We identified three possible reasons behind the poor performance of the multi-class classifier: (1) inadequate training data for different Cull classes, (2) imbalance of the training data (highly skewed towards U.S. No. 1 sweetpotatoes, and (3) similarities among the Tailed, Tapered, and U.S. No. 1 shape classes.

**Table 4.** Confusion matrix for gradient boosting algorithm evaluated on holdout data for multi-class classification. TP stands for True Positive. We can see that the model performs well in predicting U.S. No. 1 and Round sweetpotatoes, but fails to distinguish among other Cull types.

| Count | Prediction |  |  |  |  |  |
| --- | --- | --- | --- | --- | --- | --- |
| n=266<br>True Class | U.S. No. 1 | Curved | Round | Tailed | Tapered | Other |
| U.S. No. 1 (n=123) | 113 (TP) | 1 | 2 | 6 | 0 | 1 |
| Curved (n=51) | 20 | 27(TP) | 0 | 3 | 1 | 0 |
| Round (n=26) | 1 | 0 | 21(TP) | 2 | 0 | 2 |
| Tailed (n=27) | 17 | 1 | 0 | 6(TP) | 3 | 0 |
| Tapered (n=24) | 13 | 4 | 2 | 0 | 5(TP) | 0 |
| Other (n=15) | 9 | 3 | 2 | 1 | 0 | 0(TP) |

For calculating shape features, we used NIR images, which do not have any color information. Color images, which are also captured by the Exeter sorter, may provide more information with regard to certain shape defects. In future works, it would be worth investigating possible correlations between shape defects and color (e.g., defective regions may have a different color pattern than the rest of the crop). These additional color features may further improve classification accuracy, and also provide novel insight into crop quality.

## Conclusion

The irregular structure of many horticultural crops makes shape feature extraction a challenging task. We have introduced a method that extracts multiple 3D shape features from crop images captured by industrial sorters. As a first approach towards automated shape phenotyping at a large scale, our method shows promising results and the potential to be used in industrial sorters. The major contributions of our approach are 1) the capability of capturing shape features for thousands of sweetpotatoes and assess their variation across cultivars and shape classes, 2) the identification of 3D shape features that are important to determining SP shape class, and 3) the downstream use of these features to create machine learning algorithms for automated sweetpotato shape class determination. We have provided an example of the application of our method in quantifying shape variations across SP cultivars. This work opens up possibilities for creating a large scale SP shape database, which can be coupled with agricultural data to make inferences about SP shapes based on extrinsic factors (i.e., weather, cultural practices, and soil type). Importantly, the applicability of our feature extraction method is not limited to sweetpotato. This approach can be used for analyzing shapes of other vegetables and fruits (i.e., carrot, strawberry, apple) that are sorted using the Exeter sorter or a sorter with similar imaging capabilities. Machine learning classifiers for other crops can also be trained by creating crop-specific labeled datasets. One limitation of our approach is the dependency on predefined features to classify shapes. Existing deep learning methods that can extract inherent features from image data may yield higher classification accuracy. However, training such models will require a significantly large amount of labeled data to train millions of model parameters. We believe that this avenue needs to be explored in future work. With the incorporation of additional labeled images, deep neural network approaches might further increase the classification accuracy and reduce dependency on engineered features.

## Supporting information

**S1 Text Camera Scale factor calculation**

**S2 Text Sweetpotato Shape Features Calculation**

**S3 Data Sweetpotato Shape Data**

**S4 Text Hyperparameters for Classification Models**

**S5 Table Model evaluation results**

## Acknowledgments

We would like to thank SAS Institute for providing support and software access for this research, and Exeter Engineering Inc. for their support in the imaging data collection process. This work was funded in part via grant (1404) 2020-1160 awarded to CMW through North Carolina State University’s Plant Sciences Initiative (NC PSI) Game-Changing Research Incentive Program. Funding from the North Carolina SweetPotato Commission for breeding research is also acknowledged.

