## Supplementary material for "Computer vision approach to characterize size and shape phenotypes of horticultural crops using high-throughput imagery": S1 Text

### SF1. Camera Scale Factor Calculation

We used a model sweetpotato of known dimensions to calculate the camera scale factor. We scanned the model sweetpotato using the Exeter Accuvision sorter and applied our 3D reconstruction algorithm to regenerate the 3D structure (Fig 2). Straight length and width captured from the feature extraction algorithm were in an arbitrary unit (A.U.). We used the known straight length to calculate the scale factor for converting A.U. into inches.

$$Scale\ Factor = \frac{Known\ True\ Length\ (inches)}{Estimated\ Length\ (A.U.)} \quad (1)$$

We scanned the model sweetpotato *nine* times using the scanner and obtained *nine* reconstructed 3D models. Next, we averaged scale factors obtained from all the scans to estimate camera scale factors. This scale factor was used to convert all arbitrary units in the model to measurement in inches. The model sweetpotato was 8 inches long (straight length) and 3.1 inches wide (Width/ Maximum Diameter). The calculated scale factor is 0.0164 inches. Distribution of scale factors over nine separate scans is depicted in Fig 1.

After adjusting the scale, average sum squared error for straight length measurement of the model SP was 0.054 $inches^2$. Average sum squared error for width measurement was 0.2049 $inches^2$.

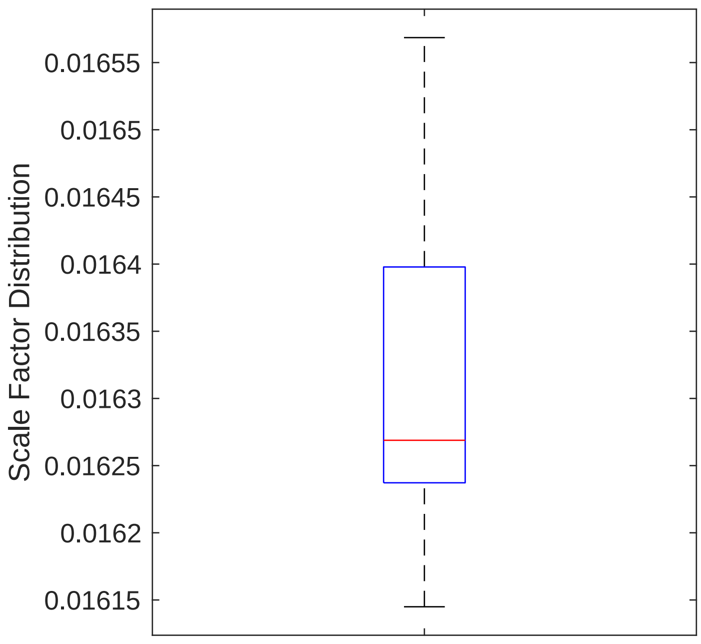


Figure 1: Distribution of scale factors from individual scans of the model sweetpotato.

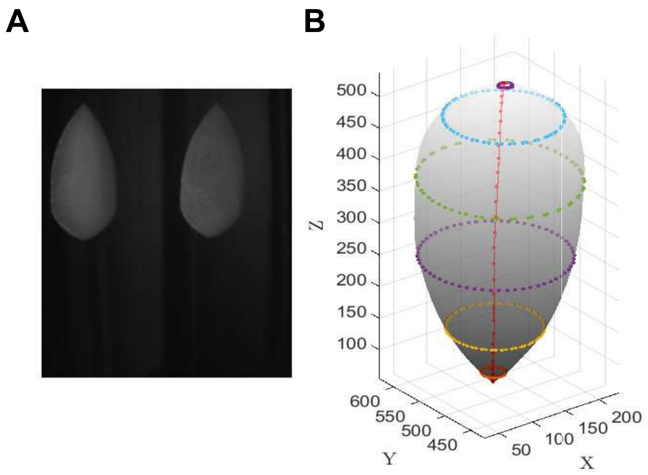


Figure 2: (A) NIR image of the model sweetpotato captured by Exeter Accuvision sorter, (B) 3D reconstruction of the model in arbitrary unit (A.U.)
