## Supplementary material for "Computer vision approach to characterize size and shape phenotypes of horticultural crops using high-throughput imagery": S2 Text

### SF2. Shape Features Calculation

Each 3D sweetpotato model was created using 20 ellipses (on the $XY$ plane) along the $Z$ axis. These ellipses were interpolated to generate an ellipsoidal 3D model of sweetpotato. We calculated 31 equidistant centroids across the $Z$ axis. Then we identified SP cross-sections that are normal to the SP curved axis for these 31 points. Each cross-section was identified by calculating the intersection between the 3D SP model and a plane normal to the curved axis at a given point (See Fig 1). Giving us 31 cross-sectional planes that are normal to the curved axis.

#### Straight Length $L_s$

Tip to tip length was estimated by measuring the distance between two furthest points on the curved axis of the 3D sweetpotato model.

$$L_s = ||C_1 - C_{31}|| \tag{1}$$

where $C_1$ and $C_{31}$ are two tips of the sweetpotato and $||.||$ is the Euclidean distance operator.

#### Curved Length $L_c$

Curved length was calculated by adding the distances between the cross-section centers along the SP axis. We discretized the curved axis at 31 equidistant points (See Fig 1 A). Curved length is calculated using the following equation

$$L_c = \sum_{i=2}^{i=31} ||\mathbf{C}i - \mathbf{C}_{i-1}|| \tag{2}$$

#### Diameter across Cross-sections

We calculated the diameter for each cross-section by finding the maximum distance between points on the SP surface (See Fig 1 B). Diameter of a cross-section $D_i$ is calculated as

$$D_i = max_{k \neq j}(||P_k - P_j||) \tag{3}$$

Where $P_k$ and $P_j$ are two points on SP surface at the $i^{th}$ cross-section.

#### Maximum Diameter / Width

We calculated the width (W) by finding the maximum cross-sectional diameter.

$$W = max_{i=1}^{i=31}(D_i) \tag{4}$$

#### Tail Length $L_{Tail}$

Tail length $L_{Tail}$ was estimated by calculating the length at two tips of the sweetpotato where the cross-section diameter is less than or equal to 0.5 inches.

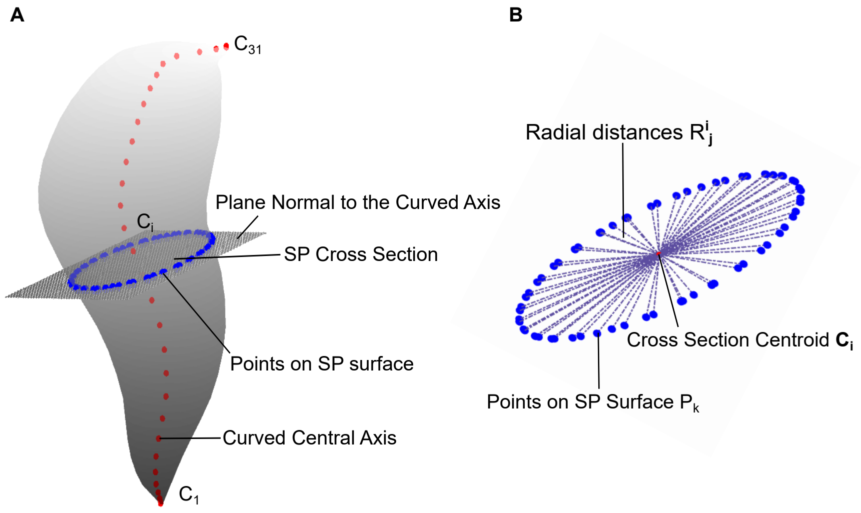


Figure 1: (A) Calculation of SP cross-sections along the curved axial length. Cross-sections are normal to the curved axis. Red dots indicate the points where cross-sections were estimated (cross-section centroid). (B) Shows an enlarged image of a cross-section. Radial distances are calculated from cross-section center to points of SP surface. Standard deviation of these radial distances are used to calculate the roundness measure of a specific cross-section.

#### Cross-Section Roundness

For the $i^{th}$ cross-section we measured radial distances as $R_j^i = ||C_i - P_k||$ (See Fig. 1B) Cross-section roundness $\sigma_{R^i}$ is the standard deviation of these radial distances normalized by their mean.

#### Average Cross-section Roundness

This feature was calculated as $\frac{\sum_{i=1}^{i=N} \sigma_{R^i}}{N}$ where N=31.

#### Volume (V)

For each cross-section we calculated the area $\mathbf{A_i}$ using the *polyarea* function of MATLAB. This function calculates the area defined by the **P** points (See Fig 1). We obtained the volume of SP by integrating the areas across the curved length (**C**).

$$V = \int_{C_1}^{C_N} \mathbf{A}.\mathbf{C} d\mathbf{C} \quad (5)$$

We applied the trapezoidal integration rule to calculate the volume (using the ***trapz*** function in MATLAB).
