## Supplementary material for "Computer vision approach to characterize size and shape phenotypes of horticultural crops using high-throughput imagery": S4 Text

### SF4.Hyperparameters for Classification Models

We obtained hyperparameters for each model using the genetic search algorithm in SAS Viya software. For all models, we used a five-fold cross-validation (70% training and 30% validation split) on the training data to tune these parameters. Optimized parameters for models are given in following tables. Details of these paremeters can be found in the SAS Viya Documentation[1]

#### 1 Binary Classification

#### Neural Network (Champion Model)

Table 1: Optimized hyperparameters for the neural network model (Binary classifier)

| Variable Name | Value |
|---|---|
| Number of Hidden Layer | 1 |
| Neurons in Hidden Layer 1 | 100 |
| Auto tuning Search Method | Genetic Algorithm |
| Optimization Method | LBFGS |
| Activation function | Hyperbolic Tangent |

#### Random Forest

Table 2: Optimized hyperparameters for the random forest model (Binary classifier)

| Variable Name | Value |
|---|---|
| Number of Trees | 150 |
| Bootstrap | 0.1 |
| Maximum Tree Levels | 16 |
| Number of Bins | 100 |

#### Decision Tree

Table 3: Optimized hyperparameters for the decision tree model (Binary classifier)

| Variable Name | Value |
|---|---|
| Maximum Tree Levels | 20 |
| Maximum Bins | 93 |
| Criterion | Gini |

[1] https://go.documentation.sas.com/?docsetId=webeditorref&docsetTarget=p04w1umaff496dn1evuvs0krzt2p.htm&docsetVersion=3.71&locale=

#### Gradient Boosting

Table 4: Optimized hyperparameters for the gradient boosting model (Binary classifier)

| Variable Name | Value |
|---|---|
| Learning Rate | 0.12 |
| Sampling Rate | 1 |
| Lasso | 6.67 |
| Ridge | 2.22 |
| Number of Bins | 96 |
| Leaf Size | 1 |

#### Bayesian Network

Table 5: Optimized hyperparameters for the Bayesian network model (Binary classifier)

| Variable Name | Value |
|---|---|
| Maximum Number of Parents | 2 |
| Parenting Algorithm | BESTSET |
| Network Structure | Tree Augmented Naive Bayes (TAN) |
| Number of Bins | 12 |

### 2 Multi-class Classification

#### Neural Network

Table 6: Optimized hyperparameters for the neural network model (Multi-class classifier)

| Variable Name | Value |
|---|---|
| Number of Hidden Layer | 1 |
| Neurons in Hidden Layer 1 | 51 |
| Auto tuning Search Method | Genetic Algorithm |
| Optimization Method | LBFGS |
| L1 Regularization | 0.001 |
| L2 Regularization | 0.001 |
| Activation function | Hyperbolic Tangent |

### Random Forest

Table 7: Optimized hyperparameters for the random forest model (Multi-class classifier)

| Variable Name | Value |
|---|---|
| Number of Trees | 85 |
| Bootstrap | 0.5 |
| Maximum Tree Levels | 30 |
| Number of Bins | 60 |

### Decision Tree

Table 8: Optimized hyperparameters for the decision tree model (Multi-class classifier)

| Variable Name | Value |
|---|---|
| Maximum Tree Levels | 16 |
| Maximum Bins | 143 |
| Criterion | CHISQUARE |

#### Gradient Boosting (Champion Model)

Table 9: Optimized hyperparameters for the gradient boosting model (Multi-class classifier)

| Variable Name | Value |
|---|---|
| Learning Rate | 0.34 |
| Sampling Rate | 0.8 |
| Lasso | 10 |
| Ridge | 6.67 |
| Number of Bins | 100 |
| Leaf Size | 12 |

#### Bayesian Network

Table 10: Optimized hyperparameters for the Bayesian network model (Multi-class classifier)

| Variable Name | Value |
|---|---|
| Maximum Number of Parents | 2 |
| Parenting Algorithm | BESTONE |
| Network Structure | NAIVE |
| Number of Bins | 5 |
